# Notch pathway is required for protection against heat-stress in spermatogonial stem cells

**DOI:** 10.1101/2021.06.10.447875

**Authors:** Omar D. Moreno Acosta, Agustín F. Boan, Ricardo S. Hattori, Juan I. Fernandino

## Abstract

Environmentally favorable conditions the sustainability of spermatogenesis is brought about by a balance between two types of division, the self-renewal division for the maintenance of the stem cell pool and the differentiation division for continuous production of spermatozoa. The production of gametes under unfavorable, stressful conditions can decrease or even be interrupted, compromising fertility parameters. Thus, the survival of spermatogonial stem cells (SSCs) is crucial for the recovery of spermatogenesis after stressful situations (e.g. high temperature). Here, we show that the Notch pathway protects the spermatogonial stem cells against thermal stress, ensuring reproductive success after normal conditions are restored. First, presenilin enhancer-2 (pen-2), the catalytic subunit of γ-secretase complex, was localized in SSCs of the medaka testis. The exposure of adult males to thermal stress condition induced apoptosis in all spermatogenics cells, with the exception of SSCs. Concomitantly, the Notch pathways was up-regulated, including the *pen-2*, its ligands (dll4, jag1-2) and its receptors (notch1a-3); *pen-2* expression was restricted to the SSCs during thermal stress. The importance of this pathway was further supported by an *ex vivo* approach, in which the inhibition of Notch activity induced a loss of SSCs. Overall, this study demonstrates that the Notch pathways activity is necessary for the protection of SSCs under chronic thermal stress.

## Introduction

Spermatogenesis is a cellular process necessary for the formation of male gametes from spermatogonial stem cells (SSCs), which proliferate synchronically and differentiate into millions of spermatozoa (SZ) or are constantly self-renewed asynchronically, like other stem cells [1, 2]. To maintain continuous spermatogenesis throughout the male reproductive life, SSCs reside in the “ niche”, a specialized microenvironment of the testes, which regulates their properties of self-renewal, pluripotency, quiescence, size, and their ability to differentiate and proliferate [3–6]. This ability of the niche to maintain SSCs homeostasis is what allows them to survive under adverse conditions.

Temperature is one of the most relevant environmental factors that have a strong effect on reproduction. This is due to the fact that it affects both the early development of the testes, as well as in adult stages the quality and quantity of gametes, thus compromising spermatogenesis and reproductive success [7–12]. In various groups of vertebrates, it has been studied how hyperthermia deteriorates the different types of germ cells [13], mainly due to apoptosis [14]. In several vertebrates and particularly in fish, it has been observed that under high temperatures only SSCs remain as a remnant, which have the ability to regain the germ line [15, 16]. This reversible characteristic that allows the germ line to overcome a stress factor, and the interaction of SSCs in the niche, made us hypothesize that the protection of germ cells must occur by a direct signaling between the niche and germ cells, which regulate this process.

In this sense, despite countless studies, it is still difficult to characterize the somatic stem niche in vertebrates [17] . Recent studies in early development of mice have been able to show a possible somatic stem niche [18], and much progress has also been made in other model organisms, like flies [5, 6]. In both cases, it was observed that one of the main signaling pathways involved is the Notch pathway. This is a highly conserved juxtacrine signaling pathway well characterized in other processes in mammals, mainly proliferative, and is composed of four receptors (Notch1-4) that interact with their ligands or others with similar structure, delta 1, 3-4, jagged1-2 [19, 20]. After binding to the ligand, in the canonical Notch signal pathway, the receptor is activated by the γ-secretase complex, mainly by the presenilin enhancer-2 (Pen-2) catalytic subunit, where proteolysis occurs within the transmembrane domain that releases the intracellular domain (Notch intracellular domain, Nicd) [21–23]. After cleavage, Nicd is translocated to the nucleus and associates with DNA-binding proteins, such as CSL (later CBF1), to activate transcription of cis target genes, such as hes1 and hes5 [24], that in general inhibit the expression of other genes [25]. Moreover, the participation of Notch signal, particularly Notch1, in the regulation of gonocytes quiescence has been well characterized in Sertoli cells [26]. Despite the numerous studies on the Notch pathway in the regulation of cell proliferation, little is known about its participation in the regulation of the proliferation of germ cells under an environmental stressor, such as temperature.

On these regards, in a previous study it was suggested that Pen-2 act as an anti-apoptotic gene, protecting germ cell gonocytes from temperature during testis differentiation in pejerrey fish [27]. This catalytic member of the γ-secretase complex has been shown to play an important function in the survival of cells, protecting them from apoptosis [28]. Selective knock-down of pen-2 in developing zebrafish embryos resulted in strong induction of the p53-dependent apoptosis cascade in whole animal [29]. Although the function of Pen-2 has been highly studied in brain, especially in Alzheimer’s disease [21, 30] and cancer [31], its participation on the gonad has not been fully elucidated, especially during high temperature exposure. As in fish, it has also been established that mammalian germ cells experience apoptosis via the p53 cascade during exposure to high temperatures [32]; however, germ cells, such as other stem cells [33], have a protective mechanism to avoid the induced apoptosis damage by entering a transient state of cell-cycle quiescence. Therefore, the germ cells must retain the appropriate information and totipotency, recovering later the reproduction [34].

Based on these antecedents, we considered to evaluate whether this cell-to-cell communication is involved in the mechanism of protection of germ cells against thermal stress.

## Materials and Methods

### Source of medaka

All experiments were performed with adults of medaka *Oryzias latipes* from hi-medaka strain (ID: MT835) supplied by the National BioResource Project (NBRP). These fish were kept under controlled laboratory conditions for this species [35]. Briefly, fish were reared at 26 ° C +/- 1 under a photoperiod of 14 light hours 8 of darkness. For the temperature stress experiment, the temperature in 10L fish tank was raised to 33 °C +/- 1 for 30 days. During the time that the experiment elapsed, the control group remained in the aforementioned initial conditions. Fish from each group had their testes removed for subsequent analysis at 3, 10, and 30 days after the beginning of the experiment. The control group was kept under normal conditions of the rearing room 25° C +/- 1 **(Figure 1E)**. Fish were handled in accordance with the Universities Federation for Animal Welfare Handbook on the Care and Management of Laboratory Animals (www.ufaw.org.uk) and internal institutional regulations.

**Figure 1:**
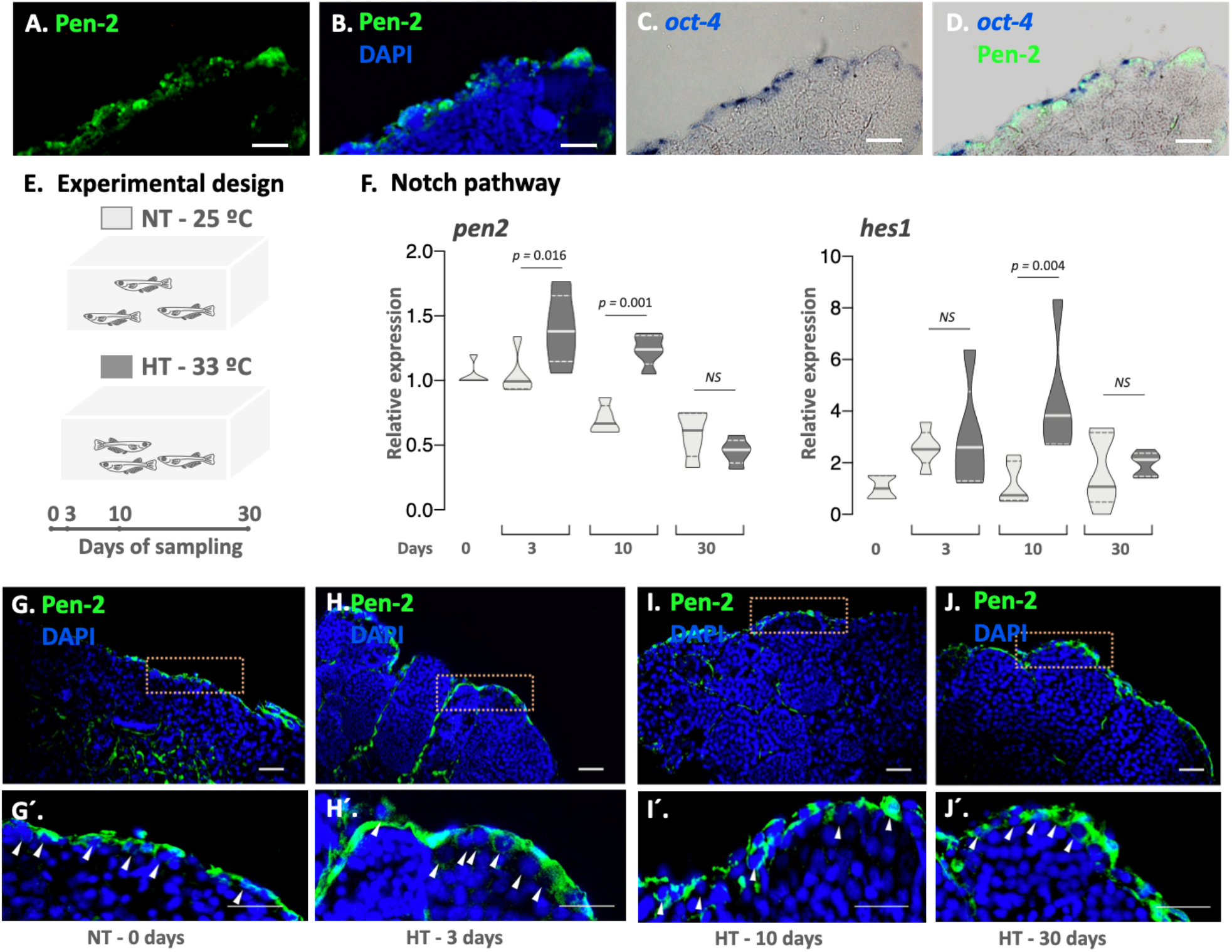
Pen-2 is down-regulated during thermal treatment in male. Transversal sections of the distal portion of the testis lobule, observing Pen-2 (green, immunofluorescence, IF) and nuclei stained with DAPI (blue) (**A, B**). Co-localization of Pen-2 (green, IF) with *oct-4* (blue, *in situ hybridization*) (**C, D**). Experimental design, in which adult males were reared at control (NT – 25°C, light grey) and high (HT – 33°C, dark grey) temperature (**E**). Notch pathway: Transcript abundance levels of *pen-2* and *hes1* in different treatment days of treatment, 0, 3, 10 and 30 days. **(F)**. IF of Pen-2 (green) in testis of male reared at HT at 0 (**G**), 3 (**H**), 10 (**I**) and 30 (**J**) days of thermal treatment, and nuclei stained with DAPI (blue). Magnification of each testis, doted orange line, at different sapling time of at 0 (**G’**), 3 (**H’**), 10 (**I’**) and 30 (**J’**) days of thermal treatment. SGa are indicate with arrowhead (**G’-J’**). Scale bar represents 20 µm. Transcript abundance quantification was performed using the 2^-ΔΔCt^ method and *pen-2* and *hes1* values were normalized to *rpl7. p*-values are indicated when transcript abundance between treatment at the same sampling day differ significantly (P<0.05). NS, not statistically significant. Relative gene expression levels were compared as described by Pfaffl [39].

### Ex vivo experiments with DAPT

To carry out ex vivo experiment with the testicular sections, the fish were anesthetized by freezing on ice and then euthanized. The testes were then dissected under the stereoscope, using forceps and sterile scissors. The gonads were removed and placed in 1 M phosphate buffer saline (PBS) with 1x streptomycin penicillin (Gibco). They were then rinsed and subsequently cut into pieces of about one millimeter. These were left overnight at 25° C, in L15 medium (Gibco) with antibiotic (Figure 4A). Then, 2 to 4 pieces of testis were placed in each well with 1 ml of medium with the DAPT drug (Selleckchem), a γ-secretase inhibitor and indirectly an inhibitor of Notch, at a concentration of 12.5 µM and 25 µM. As a control, the drug diluent was placed DMSO and incubated for 24 hours in a Fisher thermo stove.

### Total RNA Extraction and Realtime-PCR

Testes were removed from males for gene expression analysis. Total RNA extraction was carried out using 300 µL of TRIzol® Reagent (Invitrogen), according to the manufacturer’s instructions. RNA from each sample (500 ng) was used to perform the cDNA synthesis using the SuperScript II enzyme (Invitrogen).

Real-time PCR primers are listed in **Table Supplement 1**. Gene-specific qPCR was performed using the SYBR green master mix (Applied Biosystem). The amplification protocol consisted of an initial cycle of 1 min at 95°C, followed by 10 s at 95°C and 30 s at 60°C for a total 40 cycles. The subsequent quantification method was performed using the 2-ΔΔCt method (threshold cycle; assets. thermofisher.com/TFS-Assets/LSG/manuals/ cms_040980.pdf) and normalized against reference gene values for ribosomal protein L7 (*rpl7*) [36].

### Histology and immunofluorescence

Samples for histology and IF were firstly fixed in Bouin’s solution. Then they were embedded in paraffin and transversally sectioned on a Leica DM 2125RT microtome at 4-5 µm thickness.

For IF, sections were washed with 0.1 M PBS (pH 7.4) and blocked in 0.1 M PBS containing 0.5% bovine serum albumin (Sigma-Aldrich) and 0.5% Triton X100 for 60 min before overnight incubation at 4°C with primary antibody, anti-pen-2 antibody (1:200, rabbit, LS-C135520, LSBio). A negative control was also run to see specificity without primary antibody. After incubation, the sections were washed twice in PBS and incubated at RT for 90 min with Alexa Fluor 488-conjugated goat anti-rabbit IgG (ThermoFisher Scientific, A-11008) secondary antibodies at a dilution of 1:2000 in PBS. After incubation, sections were rinsed twice with PBS and mounted with Fluoromount mounting medium (Sigma-Aldrich) containing 4 ‘, 6 - diamidino-2-phenylindole (DAPI, 5 µg/ml, Life Technologies).

### TUNEL assay

The presence of apoptosis in gonad was detected through the In situ Cell Death Detection Kit, Fluorescein (Roche). Samples fixed in Bouin’s solution, embedded in paraffin and sagittal sectioned at 5 µm were treated according to the manufacturer‘s manual, with a step of permeabilization with 0.1% Triton X-100, 0.1% sodium citrate in PBS 1X solution. Fluorescein was observed under a Nikon Eclipse E600 microscope.

### RNA in situ hybridization

ISH were performed as previously described [37]. Briefly, digoxigenin-labeled probes were synthesized from the full-length medaka cDNA of *oct-4* (stem cell marker also known as pou5f1; [38]) using pGEM®-T Vector (Promega) linearized plasmid. Testicular explants from ex vivo treatment was fixed overnight in 4% RNAse-free paraformaldehyde (PFA) at 4°C, permeabilized using 20 µg/µl proteinase K at room temperature (RT), and hybridized at 68°C overnight with oct-4 digoxigenin (DIG)-labeled RNA probes. Hybridized probes were detected using an alkaline phosphatase–conjugated anti-digoxigenin antibody (1:2000; Roche) in the presence of nitro blue tetrazolium/5-bromo-4-chloro-3′-indolyphosphate substrates (Roche). Stained testicular explants were embedded in gelatin, cryostat sectioned at 14–16 µm thickness and photographed.

### Statistical analysis

Values are presented as mean ± standard error of the mean (SEM) for continuous variables and as percentages for categorical variables. Fold change and statistical analysis of RT-qPCR quantifications were performed using FgStatistics software (http://sites.google.com/site/fgStatistics/), based in the comparative gene expressions method [39]. Statistical analyses were performed by using Prism 9 (GraphPad Software, San Diego, CA). Continuous variables were compared by one-way analysis of variance (ANOVA), followed by Tukey’s multiple comparisons test, for compare the mean of each column with the mean of every other column. All differences were considered statistically significant when p < 0.05.

## Results

### Identification of Pen-2 in adult testis

The presence of Pen-2 immuno-reactive cells were observed in the distal portion of the lobule of the testis, the same germinal region of type A spermatogonia (SGa), or spermatogonial stem cell (SSCs) **(Figure 1A, B)**. To verify the co-localization of Pen-2 with SGa, an ISH with *oct-4* riboprobe was performed. I r-Pen-2 cells were observed sorrounding SGa (*oct-4* positive cells) **(Figure 1C, D)**, establishing that Pen-2 is expressed in somatic cells.

### Up-regulation of gonadal Pen-2 at high temperature

Then, we analyzed the expression of Pen-2 /*pen-2* (protein and transcript) in an *in vivo* adult treatment, keeping adult males at normal (NT-25°C) and high temperature (HT – 33°C) for 30 days **(Figure 1E)**. Firstly, we quantified the transcript abundance of Notch pathway related genes, such as pen-2 and *hes-1* (a well know Notch effector [20]). Both genes are up-regulated at HT at 3 and 10 days of treatment, showing no differences at the end of the experiment (30 days) with NT **(Figure 1F)**.

Additionally, we characterized the localization of the Pen-2 in testis of fish reared at high temperature. At the beginning of treatment (NT – 0 days) Pen-2 showed an expression in somatic cells surrounding SGa of the distal portion of the lobule, and in the medullar region of the testis, probably somatic cells surrounding spermatids in the medullar region **(Figure 1G, G’)**. Interestingly, under high temperature treatment the expression of Pen-2 is restricted to the distal portion of the lobule, where are the SGa **(Figure 1H-J, H’-J’)**.

### Apoptosis of testis germ line at high temperature

Is well know that increasing temperature induce the inhibition of spermatogenesis, with germ cell line loss. Here, high temperature during 3 days did not affect the germ line of adult males, observing spermatozoas (Sz) in the lumen of the testis **(Figure 2A)**. However, later, after 10 days of heat-treatment, we observed shorter seminiferous tubules, apparently due loss of sperm production and a large number of spermatocytes, with enlargement of the lumen. Finally, at 30 days, we observe that germ line was altered, whit low or even absent of sperm number was observed, presenting fibrosis in the medullar region. However, throughout the treatment, a large number of SSCs was observed in the testis.

**Figure 2.**
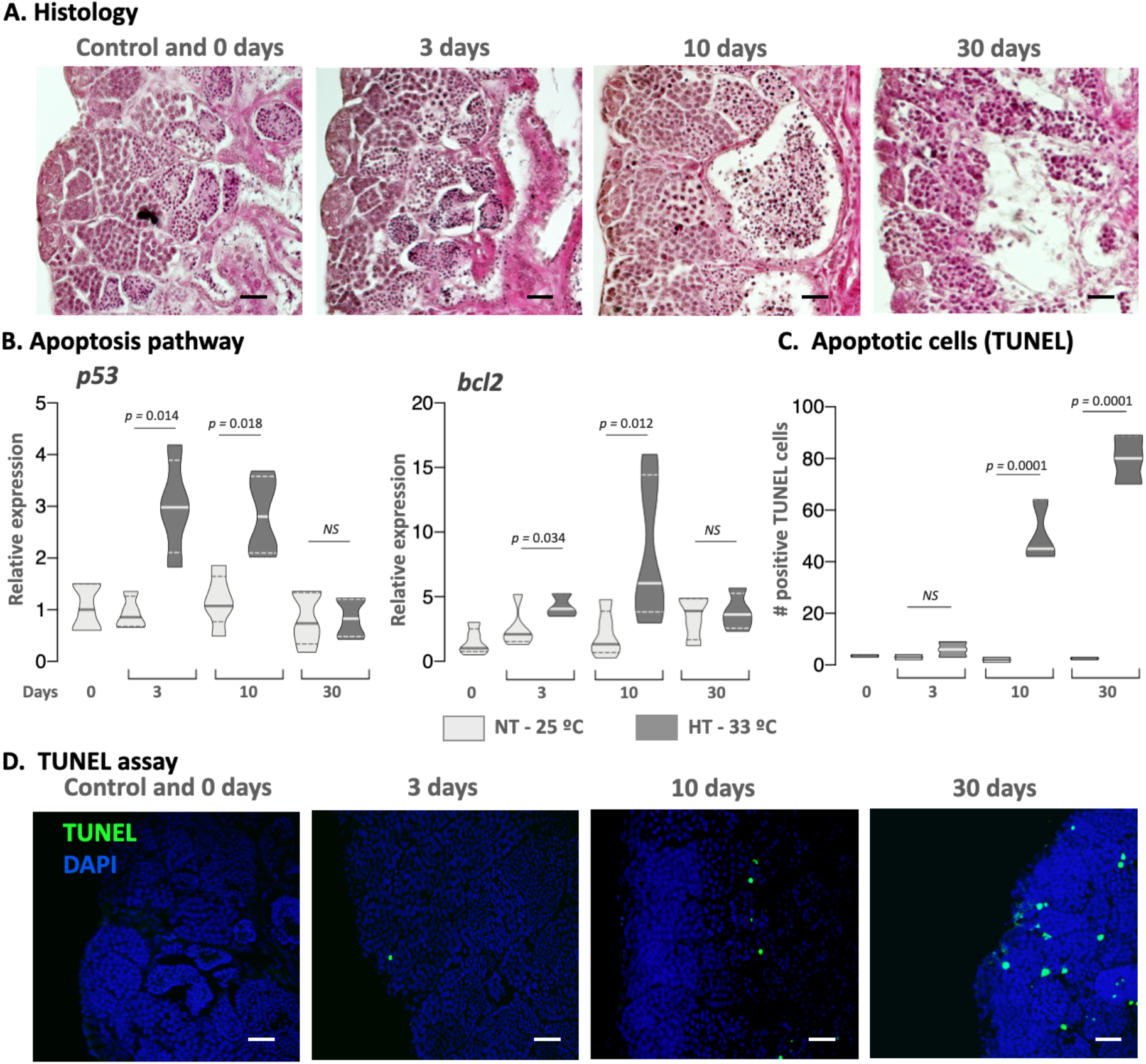
High temperature induces apoptosis of testis germ line. Histology: Transversal sections of the testis staining with eosin and hematoxylin (E&H) (**A**) of adult male reared at HT during 0 (and control), 3, 10 and 30 days. germ cell line loss. Apoptotic pathway: Quantification of *p53* and *bcl2* transcript abundance at 0, 3, 10 and 30 days in testis of males reared at normal (NT – 25°C, light grey) and high (HT – 33°C, dark grey) temperature (**B**). Apoptotic cells: Quantification of apoptotic cells by TUNEL assay at different sampling time and thermal treatment (**C**). TUNEL assay: Transversal section of the testis with TUNEL assay showing apoptotic cells (green) and nuclei stained with DAPI (blue) at 0 (and control), 3, 10 and 30 days (**D**). Scale bar represents 20 µm. Transcript abundance quantification was performed using the 2^-ΔΔCt^ method and *p53* and *bcl2* values were normalized to *rpl7*. p-values are indicated when transcript abundance between treatment at the same sampling day differ significantly (P<0.05). NS, not statistically significant. Relative gene expression levels were compared as described by Pfaffl [39] to (B); Tukey’s multiple comparisons test per (C).

To evaluate whether the temperature treatment is causing the loss of germ line, we decided to evaluate apoptosis in the heat-treatment. The apoptotic pathway, quantified by *p53* and *bcl2* gene showed the same pattern of *pen-2*, with an up-regulation in HT testis at 3 and 10 days in comparation to NT **(Figure 2B)**, establishing that the heat-treatment induced apoptosis. Finally, to corroborate the loss of germ line by high temperature we quantified apoptosis by TUNEL assay. We observed that the loss of the germ line is caused by apoptosis for 10 days of treatment, showing an increased number of positive tunnel cells **(Figure 2C, D)**. Moreover, at 30 days, the number of cells positive tunnel is higher compared to the control group. This greater number of apoptotic cells are found in the medullar region of the testis, whereas no apoptotic cells were observed in the peripheral region of the tubule, where oct4-positive cells or SGa are observed **(Figure 2D)**.

### Activation of Notch pathway in testis under high temperature

Taking into account the expression of Pen-2/**pen-2** and **hes1**, two key players in the Notch pathway, in testes kept at high temperature, we decided to measure the transcript levels of ligands and receptors of this pathway at 10 days of treatment **(Figure 3A)**. The transcript abundance of the ligands *dll4, jag1a, jag1b* and *jag2*, as well as the receptor notch-1a were up-regulated at high temperature compared to the controls **(Figure 3B)**. Otherwise, *notch3* and *jag2b* did not show significant differences **(Figure 3B)**.

**Figure 3.**
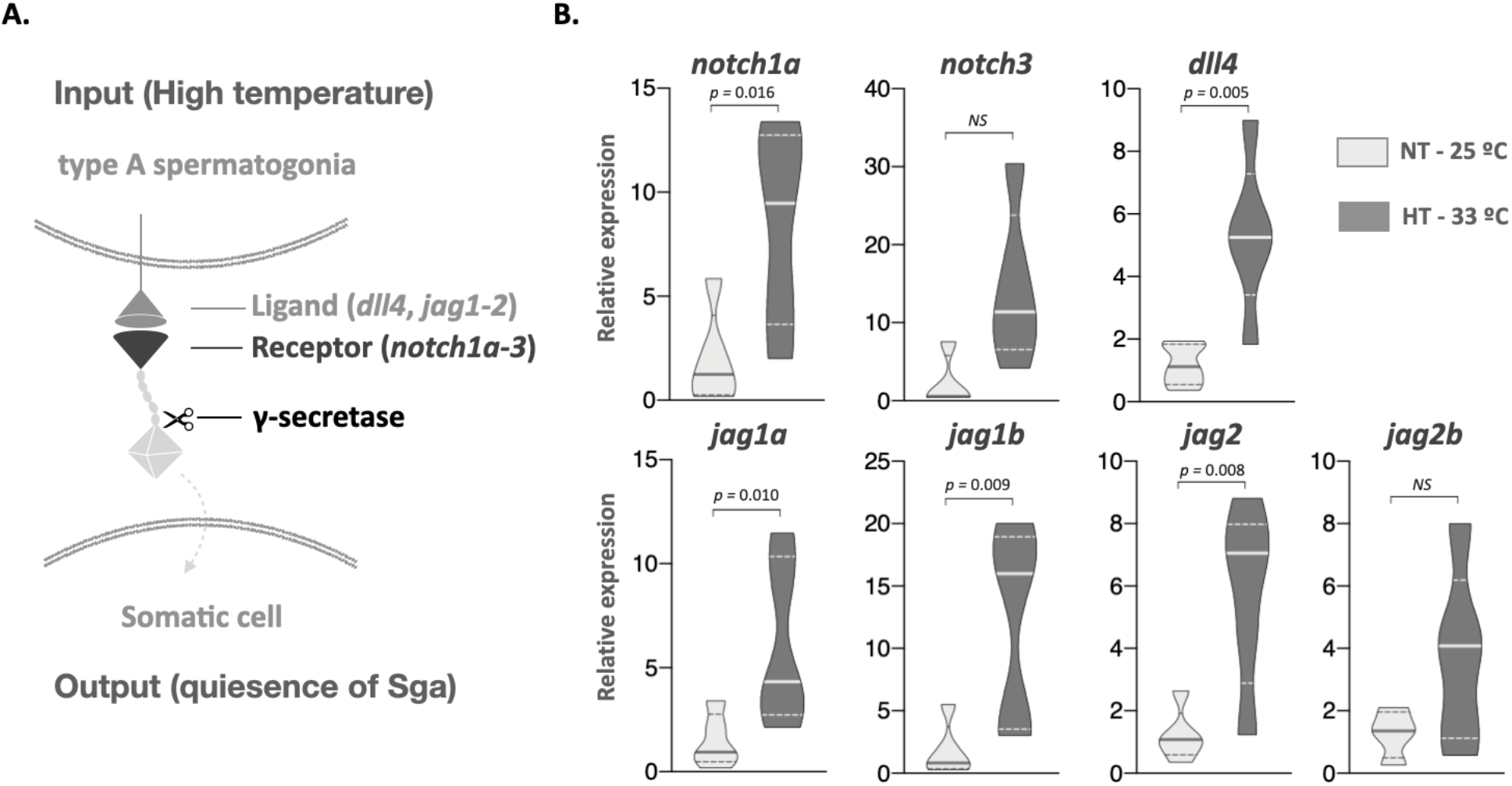
Notch pathway is up-regulated in testis at 10 days of heat treatment. Presumptive Notch cell-to-cell signaling between type A spermatogonia and Sertoli cell under a high temperature input (modified from Henrique & Schweisguth [20]) (**A**). Quantification of different Notch pathway ligand (*dll4, jag1a-b, jag2-2b*) and receptors (*notch1a, 3*) transcript abundance at 10 days in testis of males reared at normal (NT – 25°C, light grey) and high (HT – 33°C, dark grey) temperature (**B**). Transcript abundance quantification was performed using the 2^-ΔΔCt^ method and values were normalized to *rpl7*. p-values are indicated when transcript abundance between treatment at the same sampling day differ significantly (P<0.05). NS, not statistically significant. Relative gene expression levels were compared as described by Pfaffl [39].

**Figure 4.**
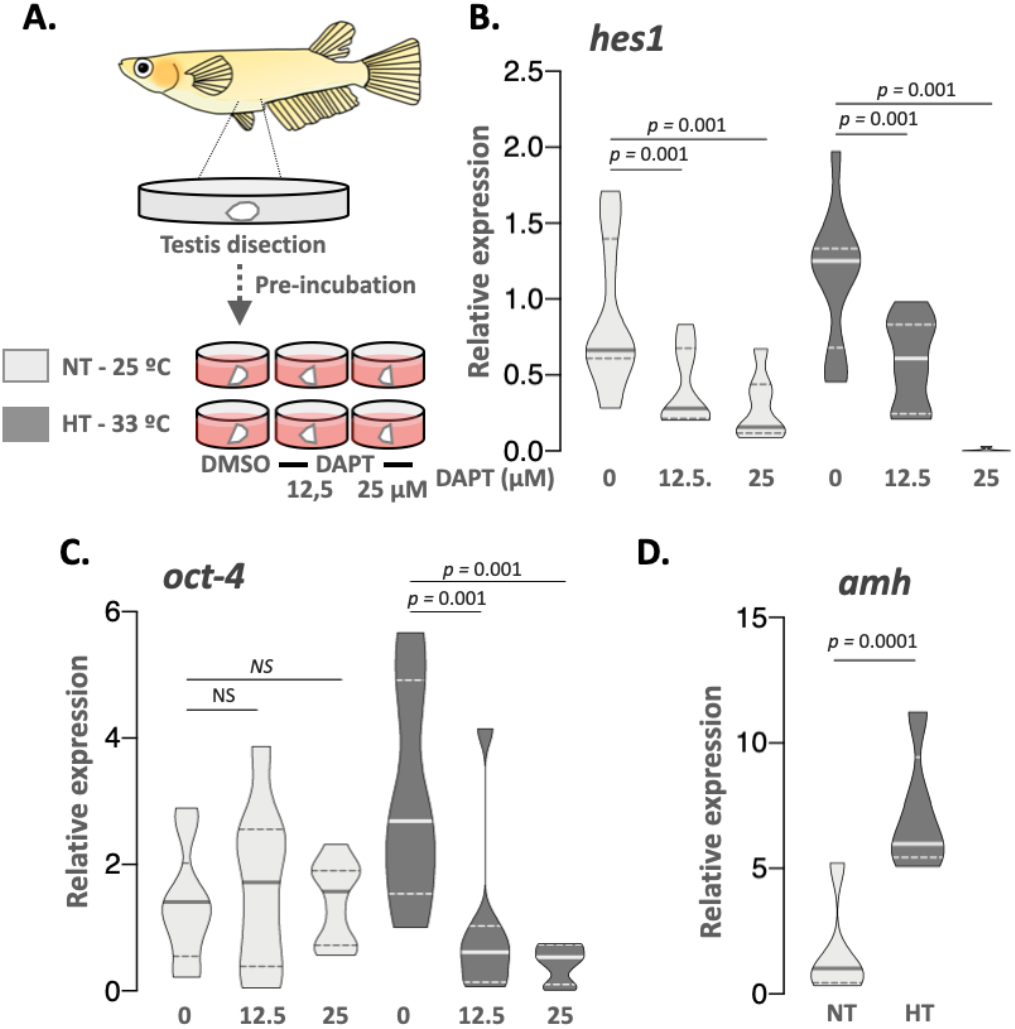
Notch pathway protects type A spermatogonia to heat-induced loss. Experimental design of *ex vivo* approach, in which adult testis explants were incubated during 24 hours at control (NT – 25°C, light grey) and high (HT – 33°C, dark grey) temperature with DAPT (γ-secretase inhibitor; 12,5 and 25 µM) by triplicated (**A**). Transcript abundance levels of *hes1* (**B**, Notch pathway cis target genes) and *oct-4* (**C**, stem state marker gene) to different treatments. Transcript abundance of *amh* of testis explant incubated at NT and HT during 24 hours (**D**, germ cell proliferation inhibitor). Transcript abundance quantification was performed using the 2^-ΔΔCt^ method and *pen-2* and *hes1* values were normalized to *rpl7*. p-values are indicated when transcript abundance between treatment at the same sampling day differ significantly (P<0.05). NS, not statistically significant. Relative gene expression levels were compared as described by Pfaffl [39].

### Inhibition of Notch pathway in ex vivo testis explants at high temperature

After observed an up-regulation of Notch pathway with restriction of Pen-2 in the SGa region of the distal lobule, our next step was inhibited this pathway under high temperature treatment. Given the complexity of the Notch pathway and how it affects the fate of many cell types in different tissues, we decided to use an ex vivo testis explants approach to analyze the protective role of this cell-to-cell signaling. Initially, we detected that apoptosis was induced from 3 hours of exposure, observing the up-regulation of p53 **(Figure Supplement 1A)**. Moreover, no differences were observed between 3, 12 and 24 hours of high temperature exposure. Additionally, dose-response curve showed that 12.5 (moderate down-regulation of *hes1*) and 25 µM (high inhibition of *hes1*) of the Notch inhibitor DAPT were the best condition to block the Notch action **(Figure Supplement 1B)**.

Then, we observed that *hes1* was down-regulated in a dose-dependent manner, at both normal and high temperature when ex vivo testis explants were treated with DAPT **(Figure 4B)**, showing that Notch pathway was inhibited. Interestingly, the expression of the stem state marker gene *oct-4* was affected only by temperature treatment **(Figure 4C)**, suggesting a reduction in the SGa and a necessary protective effect of Notch on germ cell fate under thermal stress.

In the next step the protective effects of Notch pathway at high temperature exposure in avoiding the loss of SGa after 24 hours of incubation was analyzed. Since no apoptosis in SGa was observed in the *in vivo* temperature experiment **(Figure 2A)**, we studied SGa quiescence by means of the levels of the Amh, a germ cell proliferation inhibitor [40].

We observed that the levels of *amh* increase significantly when the testes are subjected to heat stress **(Figure 4D)**, suggesting that the increase in temperature promotes the quiescence of the germ cells through the activation of the Notch pathway.

## Discussion

In the last decades a decrease in the seminal parameters of men has been observed, mainly attributed to changes in behavior, as well as to increases in temperature [41, 42], which is a well -known stressor that disrupts spermatogenesis even in fish. Once the stressful condition is overridden, the recovery of spermatogenesis is a necessary mechanism to ensure the continuity of reproduction of an individual. This can be achieved because spermatogonia, which are stem cells, shift to a quiescence state in order to protect themselves from the harmful environment [43]. Despite the reproductive importance of recovery of spermatogenesis after stress, the molecular regulators that protects these germ stem cells against an increase in temperature have not been fully elucidated. In the present study, we demonstrated that exposure to high temperature in adult medaka males induce the activation of the juxtracrine signaling system of Notch in somatic cells, presumably Sertoli cells [44], surrounding type A spermatogonia. The inhibition of this pathway under thermal stress produced a fast loss of germ stem cells, supporting that the activation of the Notch pathway is essential for the maintenance of the germ line, ensuring the re-establishment of spermatogenic cycle.

Although increasing temperature promotes spermatogenesis by differentiation of spermatogonia [45], prolonged persistence at high temperatures causes reduction in the spermatogenic cell line due to an increase in apoptosis in most cells [46, 47], with the exception of type A spermatogonia [16, 47, 48]. These stem cells are kept in a quiescent state, a strategy that protects them from stress by inhibiting differentiation-activating signals, either intrinsic or mediated by cell-to-cell interactions [49–51]. For this reason, the interpretation of the mechanism that allows these cells to persist during stress would have important implications for understanding the reduction of male fertility, or even infertility. In our experimental model medaka, thermal stress showed to induce the loss of spermatogenic line, from spermatocytes to spermatozoa. This apoptotic induction was earlier observed by the up-regulation of apoptotic-related genes, such as *p53* and *bcl2*, indicating that both the apoptotic and anti-apoptotic pathways are active under temperature stress. Apoptotic cells were observed in the medullary region of the testis, but not in the distal portion of the lobule, where spermatogonia are localized, establishing the protection of this cell type.

SSCs reside within the niche, a specialized microenvironment essential for their maintenance and self-renewal [2, 44, 52–54]. In medaka testis, the main components of the niche include SSCs directly surrounded by Sertoli cells that express sox9b [44, 55]. In the present work we observed the expression of the Notch pathway member Pen-2, a protein of the γ-secretase complex that is essential for the correct cleavage and signal transduction inside the cell [23]. Pen-2 mRNAs were restricted to the distal portion of the lobule during a thermal stress in presumptive Sertoli cells surrounding SSCs. Interestingly, Notch pathway has been observed to protect different cells from apoptosis [56]. On this regard, our observation would establish a cell-to-cell communication that protects the SSC from high temperature-induced apoptosis, with the Notch pathway holding an important role in such protective mechanism.

Apart from germ cell protection, the Notch signaling has been related to the regulation of self-renewing state and/or prevention of differentiation of SSCs through the negative regulation of glial cell line-derived neurotrophic factor (Gdnf), that is expressed in Sertoli cells [57, 58]. Gdnf has been implicated in the maintenance of the self-renewing state and/or prevention of differentiation of SSC thought the regulation of Nanos2, Stra8, Bcl6b, Cxcr4 and others in A^undiff^ spermatogonia [59–61]. Notch negative regulation of Gdnf occurs through the activation of germ cell-expressed Jag1 (jagged 1) [53, 57, 58], a ligand that was highly expressed in our thermal treatment using medaka. In case of the receptor NOTCH1, it is expressed in undifferentiated germ cells and Sertoli cells of mouse testis whereas the ligand Dll4 is ubiquitously expressed in germ cells and in some Sertoli cells [62]. During embryogenesis, the constitutive activation of NOTCH1 signaling in Sertoli cells caused the exit of gonocyte from the quiescence status [26]. In flies, Notch signaling directly controls germline stem cell development and maintenance [5, 63], reinforcing the importance of cell-to-cell Notch signaling in the regulation of germ cell differentiation. Interestingly, although our results in medaka support the activation of the Notch pathway, including receptors, ligands, γ-secretase and the respective effectors of this pathway, the activation of Notch signals seems to work in the opposite direction to that described above, wherein the input signal (high temperature) seems to be sensed by SGa while the output seems to be in the somatic cells with a high expression of Pen-2, presumably Sertoli cells [44]. This differential activation by temperature in medaka would promote the protection through a state of quiescence and the concomitant inhibition of SGa differentiation.

## Credit author statement

ODMA: Conceptualization, Methodology, Analysis; AFB: Methodology and Visualization; RSH: Writing-Reviewing and Editing, JIF: Funding acquisition, Conceptualization, Investigation, Supervision; Writing-Original draft preparation, Writing-Reviewing and Editing.

## Acknowledgements

We thank Tech. Gabriela C. López for helping with histological preparations. We also thank Dr. Tania Rodriguez for technical support and Tech. Javier Herdman (INTECH) for fish handling. We are grateful to NBRP Medaka (https://shigen.nig.ac.jp/medaka/) for providing HNI (Strain ID: MT835).

## Supplementary Files

**Figure Supplement 1.**
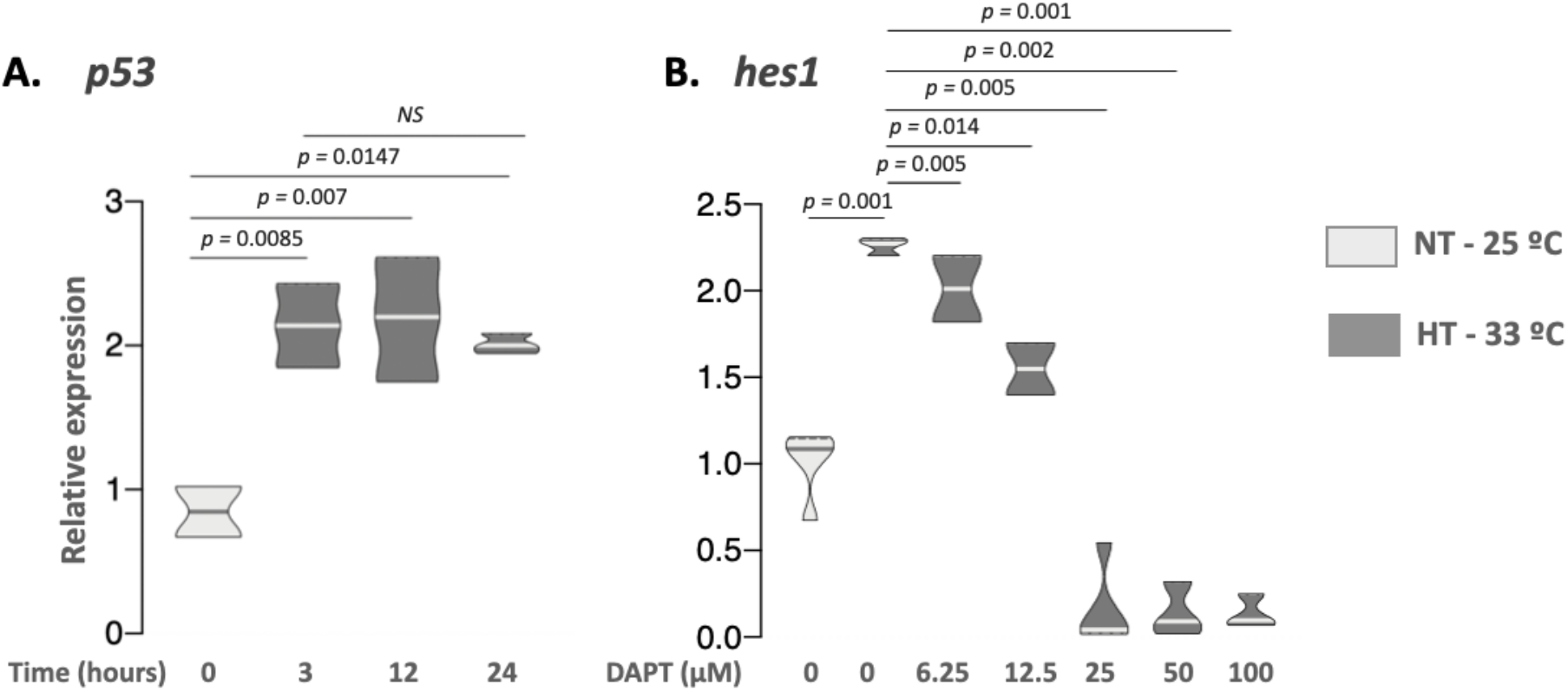
Time and doses optimization to ex vivo experiment. Time exposure curve of p53 transcript abundance in testis explants incubated at control (NT – 25°C, light grey) and high (HT – 33°C, dark grey) temperature and sampled at 0, 3, 12 and 24 hours (A). DAPT doses response: Transcript abundance levels of hes1 (Notch pathway cis target genes) in adult testis explants incubated during 24 hours at 0, 6.25, 12.5, 25, 50 and 100 µM of DAPT (γ-secretase inhibitor) (B). Transcript abundance quantification was performed using the 2-ΔΔCt method and p53 and hes1 values were normalized to rpl7. p-values are indicated when transcript abundance between treatment at the same sampling day differ significantly (P<0.05). NS, not statistically significant. Relative gene expression levels were compared as described by Pfaffl [39].

**Table Supplement 1.**
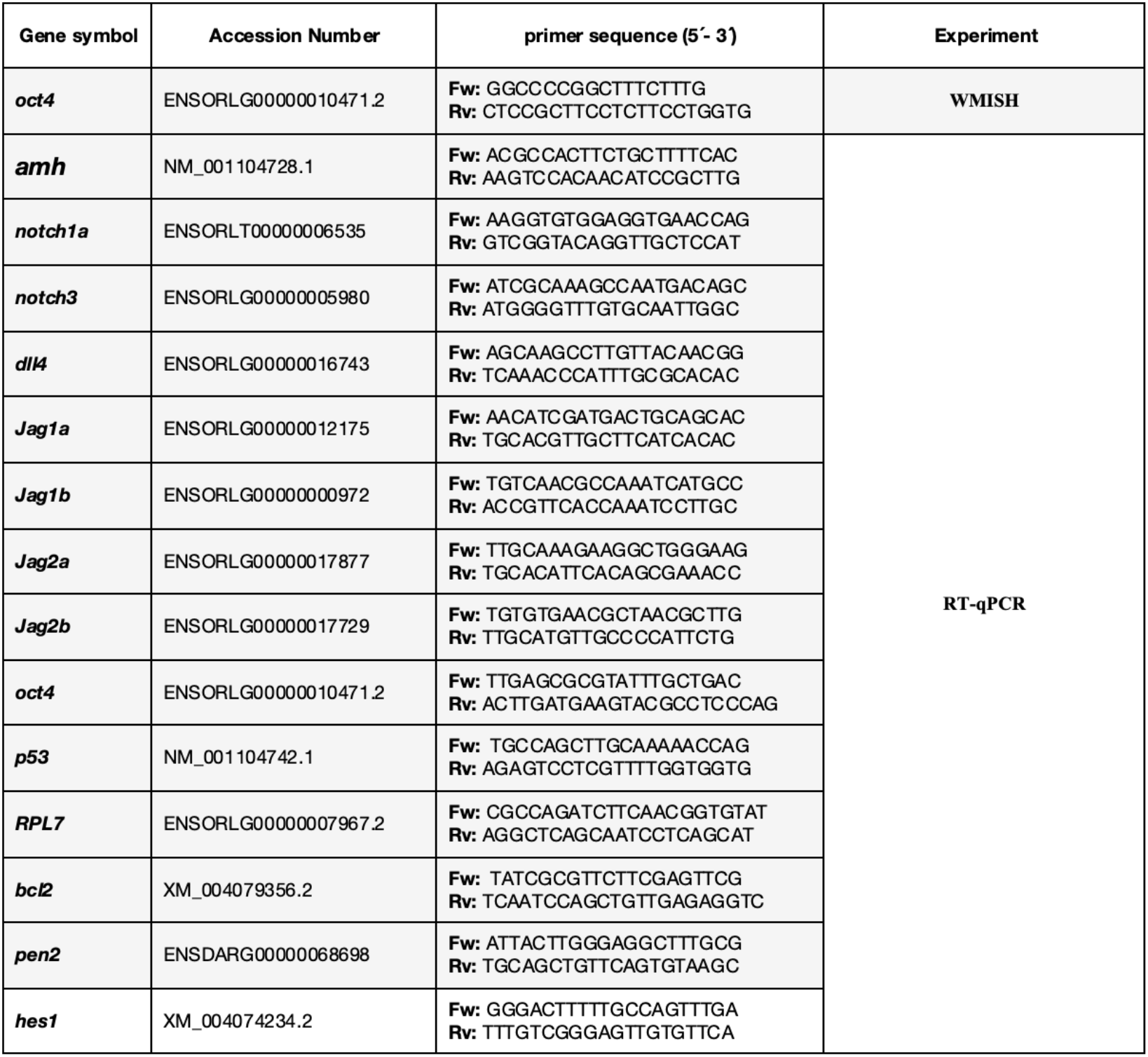
Primers sequences, ENSEMBL and NCBI accession numbers and respective references of each gene were added.

